# Characterization of the radiation desiccation response regulon of the radioresistant bacterium *Deinococcus radiodurans* by integrative genomic analyses

**DOI:** 10.1101/2021.07.07.451423

**Authors:** Nicolas Eugénie, Yvan Zivanovic, Gaelle Lelandais, Geneviève Coste, Claire Bouthier de la Tour, Esma Bentchikou, Pascale Servant, Fabrice Confalonieri

**Affiliations:** Université Paris-Saclay, CEA, CNRS, Institute for Integrative Biology of the Cell (I2BC], 91198, Gif-sur-Yvette, France

## Abstract

Numerous genes are overexpressed in the radioresistant bacterium *Deinococcus* radiodurans after exposure to radiation or prolonged desiccation. The DdrO and IrrE proteins play a major role in regulating the expression of approximately predicted twenty of these genes. The transcriptional repressor DdrO blocks the expression of these genes under normal growth conditions. After exposure to genotoxic agents, the IrrE metalloprotease cleaves DdrO and relieves gene repression. Bioinformatic analyzes showed that this mechanism seems to be conserved in several species of *Deinococcus*, but many questions remain as such the number of genes regulated by DdrO. Here, by RNA-seq and CHiP-seq assays performed at a genome-wide scale coupled with bioinformatic analyses, we show that, the DdrO regulon in *D. radiodurans* includes many other genes than those previously described. These results thus pave the way to better understand the radioresistance mechanisms encoded by this bacterium.

**Author Summary:** The main response pathway to genotoxic conditions in the radioresistant bacterium *Deinococcus radiodurans* is regulated by the constitutively expressed metalloprotease IrrE that cleaves the transcriptional repressor DdrO, leading to the expression of the genes repressed by DdrO. One of the major goals to better understand how pathways involved in radioresistance are coordinated into this fascinating bacterium is to highlight genes regulated by DdrO. In this study, we mapped *in vivo* the DdrO regulon in *D. radiodurans* by using two genome-scale approaches, ChIP-seq and RNA-seq analyses, coupled with bioinformatic analyses. As homologs of these two proteins are also found in many other bacteria, these results also pave the way to compare the stress-induced responses mediated by this couple of proteins in diverse bacteria.

## Introduction

*Deinococcus radiodurans* is one of the most resistant bacteria to genotoxic agent exposure and desiccation isolated so far [1–4]. Unlike radiosensitive organisms, once exposed to huge γ-rays doses, or after prolonged desiccation, *D. radiodurans* is able to reconstruct an intact genome in a few hours from several hundred DNA fragments [5]. Many factors contribute to the radioresistance of *D. radiodurans,* including efficient DNA repair mechanisms [5–8], a condensed nucleoid limiting the dispersion of genome fragments after irradiation [9, 10] and the protection of proteins against oxidative damage [11]. Thus, the exceptional ability of this bacterium to overcome severe DNA damaging conditions is described as a combination of active and passive mechanisms acting in synergy within the cell enabling survival from these stresses.

*D. radiodurans* exposure to γ-rays, or after recovering from desiccation, results in a rapid upregulation of the expression of many genes, including several DNA repair genes [12, 13]. In many bacterial species, expression of DNA repair genes is under the control of LexA, the repressor of the well-known SOS response (for review [14]). *D. radiodurans* encodes two LexA homologs (LexA1 and LexA2) that undergo, as in *E. coli*, a RecA-dependent cleavage after DNA damage, but neither LexA1 nor LexA2 are involved in the induced expression of RecA (15,16]. In *Deinococcus*, the main response pathway to genotoxic conditions is regulated by the constitutively expressed metalloprotease IrrE [17, 18] and the transcriptional repressor DdrO [19, 20]. *In vivo*, the loss of function of IrrE completely abolished the induction of expression of numerous genes after exposure to ionizing radiation and resulted in significant sensitivity of the strain to genotoxic conditions [19–25]. When cells are exposed to genotoxic stress conditions IrrE stimulated by an increased availability of zinc ions, cleaves the C-terminal region of DdrO [17,25,26], abolishing its DNA binding properties and leading to the expression of the genes repressed by DdrO [19].

The DdrO protein is composed of two domains: an N-terminal helix-turn-helix (HTH)_XRE DNA-binding domain itself associated to a specific structural domain at the C-terminus required for protein dimerization and for DNA binding [27, 28]. *In vitro*, IrrE-mediated cleavage removes the C-terminal 23 amino acid residues from DdrO [17, 27].

The *ddrO* gene is essential for cell viability of *D. radiodurans* and *Deinococcus deserti* [17, 20]. Interestingly, its prolonged depletion by a conditional deletion system induces, in *D. radiodurans*, an apoptotic-like response (DNA degradation, defects in chromosome segregation, membrane blebbing) [20]. These results suggested that, as in eukaryotic cells, management of DNA damage can lead to cell survival or cell death [29]. In *D. radiodurans* these two responses are mediated by common regulators, IrrE and DdrO.

The IrrE/DdrO protein pair is highly conserved in *Deinococcus* species and genes encoding IrrE/DdrO-like proteins are also present in other bacteria [30, 31]. However, questions remain about the number of the genes that are directly or indirectly regulated by these two proteins. A 17-base-pair palindromic motif, designated as the <u>R</u>adiation/<u>D</u>esiccation <u>R</u>esponse <u>M</u>otif (RDRM), was identified in the promoter regions of several radiation-induced genes in different *Deinococcus* species, suggesting the existence of a conserved radiation/desiccation response (RDR) regulon [19,32,33]. The predicted RDR regulon of 7 *Deinococcus* species consists of at least 14–24 genes, including numerous genes involved in DNA metabolism like *recA*, *ssb*, *gyrA*, *gyrB*, *uvrA*, *uvrB*, but also *Deinococcus* specific genes like *ddrA*, *ddrB*, *ddrC*, *ddrD* and *pprA* [19]. Based on the presence of the RDRM located in the promoter region of the most highly up-regulated genes by ionizing radiation and desiccation, 25 genes were predicted to belonging to the RDR regulon in *D. radiodurans*: 24 genes from Makarova et al., 2007 [33] and *ddrC* [21]. It has been shown that, *in vitro*, *D. radiodurans* DdrO was able to bind to 21 predicted RDRM motifs [26] and *in vivo*, mutations within the RDRM sequence, as well as transient depletion of DdrO, induced the expression of several RDR regulon genes [20,21,34].

In this study, we mapped the DdrO regulon in *D. radiodurans* by using two genome-scale approaches, ChIP-seq and RNA-seq analyses. These approaches were performed to find additional DdrO target sites that were not predicted by previous *in silico* analyses. To our knowledge, we present here the first ChIP-seq analysis performed at the genome level in *D. radiodurans*. Alignments of DNA sequences extracted from ChIP-seq analysis were also performed to compare the consensus motif found in the predicted RDRM. As a prerequisite to a robust feature-mapping study, we resequenced our laboratory stock strain of *D. radiodurans* R1 ATCC 13939 in order to obtain an accurate reference for read alignments and gene expression quantifications.

Our results show that the RDR regulon in *D. radiodurans* is more complex than previously thought and is composed of at least 35 genes, including genes encoding DNA and RNA metabolism proteins such as RecG and HelD helicases, and the prokaryotic replicative DNA ligase LigA, but also new genes associated with different metabolic pathways, involved in translation or encoding proteins of unknown function.

## RESULTS

### Genome sequencing

Two genome sequences of the *D. radiodurans* strain R1 are available in databases [35, 36]. The genome is distributed over 4 replicons: two chromosomes, one megaplasmid and a plasmid (Table 1). A nucleotide polymorphism between the two complete genome sequences was reported, as well as several insertions, deletions or substitutions frequently found in bacterial genomes [35]. In order to promote accurate RNA-seq and ChIP-seq analysis as well as for searching conserved binding motifs for the DdrO protein, we sequenced the *D. radiodurans* genome of strain R1 ATCC 13939 maintained in our laboratory. We opted for Illumina NextSeq Oxford sequencing coupled with Nanopore Technologies GridION to unambiguously locate the repeated elements which may misassemble short sequences in size. Merging both sets of sequences produced an ensemble of 4 high quality contigs, totaling 3578820 nt, with a 450-fold average coverage. Among of 3,230 predicted genes, 3,147 encode proteins.

**Table 1:** Size of each replicon, % GC and CDS content in Deinococcus radiodurans strain R1 ATCC 13939 genome sequence. Percentage of pairs of orthologs found in the genome sequence of our strain when compared to the two other strains R1 genome sequence when a threshold of 90 % of maximum bitscore was applied.

|  | Replicon | ID | Percentage of pairs of orthologs* | CDS | Total CDS | nt. | GC % |
| --- | --- | --- | --- | --- | --- | --- | --- |
| Hua & Hua<br>(2016) [35] | Chr 1 | CP015081 | 96,43 | 2523 | 3079 | 2646742 | 67,07 |
|  | Chr 2 | CP015082 | 96,02 | 352 |  | 433133 | 66,77 |
|  | Megaplasmid | CP015083 | 81,05 | 153 |  | 203183 | 62,98 |
|  | Plasmid | CP015084 | 62,75 | 51 |  | 61707 | 56,55 |
| White et al,<br>(1999) [36] | Chr 1 | DRA1 | 84,37 | 2629 | 3181 | 2648638 | 67,01 |
|  | Chr 2 | DRA2 | 85,05 | 368 |  | 412348 | 66,69 |
|  | Megaplasmid | DRA3 | 77,24 | 145 |  | 177466 | 63,19 |
|  | Plasmid | DRA4 | 61,54 | 39 |  | 45704 | 56,15 |
| This work | Chr 1 | DRO | 100 | 2594 | 3147 | 2644251 | 67,08 |
|  | Chr 2 | DRO_A | 100 | 364 |  | 412138 | 66,65 |
|  | Megaplasmid | DRO_B | 100 | 148 |  | 177322 | 63,21 |
|  | Plasmid | DRO_C | 100 | 41 |  | 45508 | 56,26 |

\* Percentage of pairs of orthologs (at a threshold of 90 % max bit score) found in the genome sequence of our strain when compared to each replicon from the two other strains R1 ATC 13989 genome sequence releases.

As shown in Table 1 the size of chromosome 2 and the two plasmids, as well as the number of CDS encoded by *D. radiodurans* deduced from our sequence, are closer to those published by White et al [36] than these published by Hua [35]. The large sequence insertions revealed in the more recent release were not found here. However, since the sequence published by White et al [36] contains many errors, the degree of identity of genes was better with the genome sequence published by Hua [35], with a higher percentage of genes found between these two releases when a threshold of 90 % of maximum bit score was applied (Table 1). The sequence origin for each chromosomal element and plasmids was adjusted to correspond to the genome sequence of White et al [36] and the paralogous genes mainly composed of different transposon families as well as the orthologs of each CDS, with both previous genome sequence releases listed in S1 Table.

### *In vivo* identification of DdrO binding sites by ChIP-seq assays

In order to localize *in vivo* the chromosomal regions bound by the DdrO protein, we constructed the GY 18218 strain expressing a V5-tagged DdrO protein, in all the genome copies, from the native promoter of *ddrO* (S1 Fig). Cells expressing the recombinant protein displayed the same growth rate as the wild-type strain and the expression of DdrO-V5 did not affect the resistance of the strain to DNA damaging agents (mitomycin C and UV) (S1 Fig). These results demonstrate that DdrO-V5 protein is functional and remained cleavable by IrrE under stress conditions.

*D. radiodurans* GY 18218 and R1 strains were grown to mid-log phase and ChIP-seq was performed on DNA precipitated by ChIP grade anti-V5 antibody. The Input sample (chromosomal DNA of the GY 18218 strain), the Mock sample (immuno-precipitated (IP) DNA of the wild-type strain) and three replicates of the DdrO-V5 IP sample were used to prepare sequencing libraries. The DNA regions over represented in the DdrO-V5 IP sample and corresponding to potential binding sites for DdrO-V5, were identified using the bPeaks program [37].

A total of 136 ChIP-enriched peaks were found, mainly (110/136) within intergenic regions of genes encoding proteins (Fig 1A and S2 Table), whereas 26 peaks were intragenic or found at the vicinity of genes coding for tRNA. Significant peaks, as illustrated for five genes, *ddrA, ddrB, ddrC, gyrA, gyrB* (Fig 2) were identified in the promoter region of 18 out of 25 genes reported as belonging to the RDR regulon [21,26,33]. A careful inspection of DdrO-V5 IP tag density through the IGV program of the 7 missing genes showed that a lower coverage density region was observed at the promoter region of *mutL* and small peaks were observed in the *recQ* and *sbcD* promoters that fell below the threshold used for peak detection with the bPeaks program. No peaks were detected in the region upstream of the *hutU, irrI*, *frne* and *rsr* genes.

**Fig 1:**
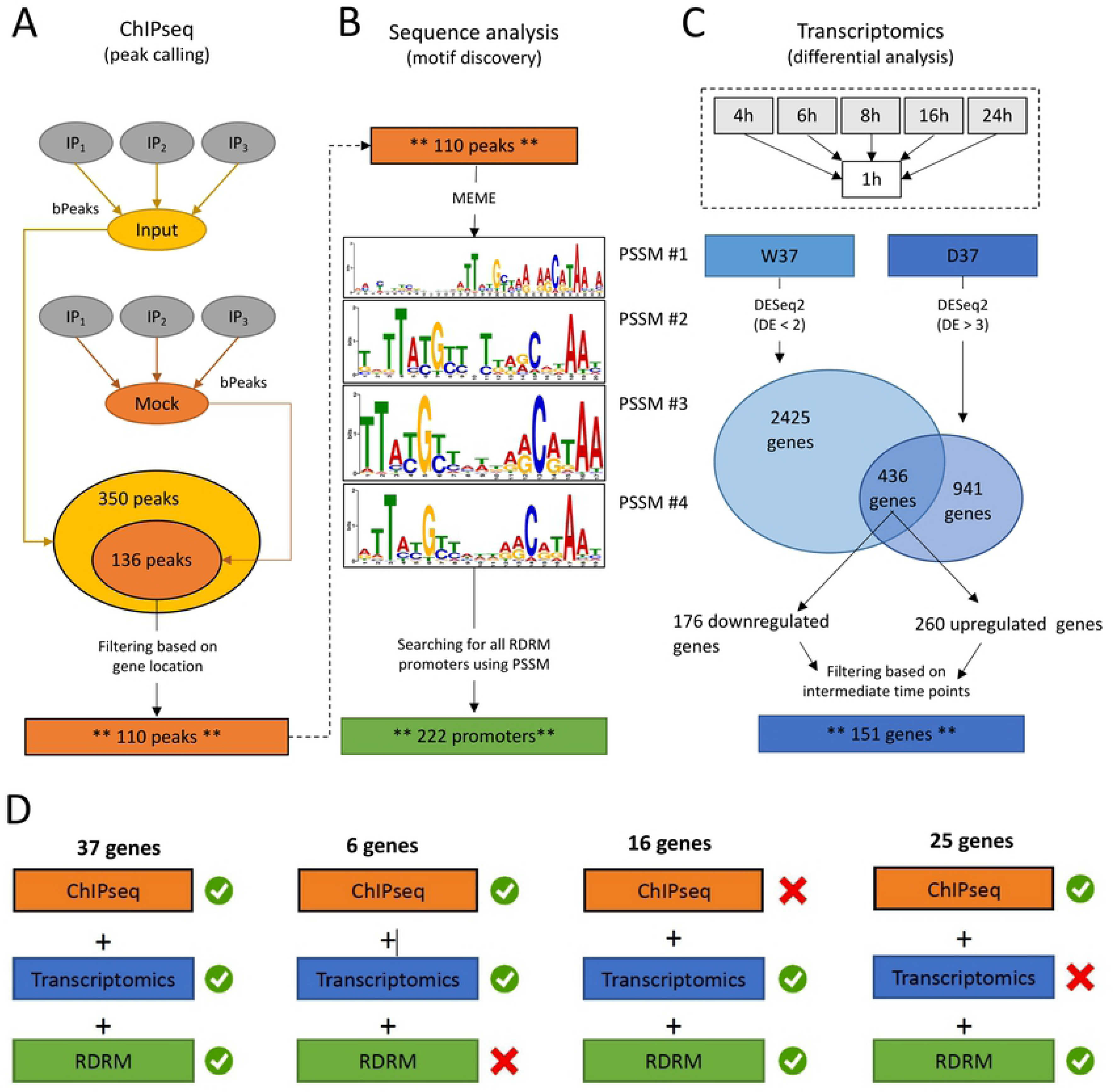
Overview of the computational strategy used to integrate omics data and identified candidate genes for inclusion in the DdrO regulon. (A). ChIP-seq analyses, *i.e.* defining the genomic regions for which interactions between DNA and the DdrO protein were observed. The three IP replicates were compared to the INPUT and MOCK controls. Peaks identified in both comparisons were retained. An additional filter was applied to focus on only the peaks located in intergenic regions. (B). Sequences of the peaks identified in (A) were used to search for over-represented DNA motifs, applying the MEME program. Four position specific scoring matrix (PSSM) were retained, because of their redundancies. PSSM were used as inputs for the FIMO program, scanning sequences between -500 and +100 of all annotated CDS. Positive matches were retained and are referred to as “RDRM promoters”. (C). RNA-seq data was used to identify differentially expressed genes, comparing each time point (4 h, 6 h, 8 h, 16 h and 24 h) to the first one (1 h). In the W37 strain, genes identified as differentially expressed in less than two comparisons were selected (DE < 2), whereas in the D37 strain, genes identified as differentially expressed in more than three comparisons were selected (DE > 3). Common genes from the two lists were retained and an additional filter was applied to focus further analyses on only these genes which exhibited differential expression (up- or down-regulation) at intermediate times, *i.e.* 6 h, 8 h and 16 h. (D). Results obtained in (A), (B), and (C) were integrated to define four levels in the DdrO regulon. The first level is comprised of genes for which (i) a peak was detected upstream of the gene location (ChIP-seq results), (ii) specific differential expression was observed in the D37 strain (transcriptome results) and (iii) an RDRM motif was found in the gene’s promoter (sequence analysis). Other levels matched with only two criteria. These genes are worth considering as being potential targets for DdrO.

**Fig 2:**
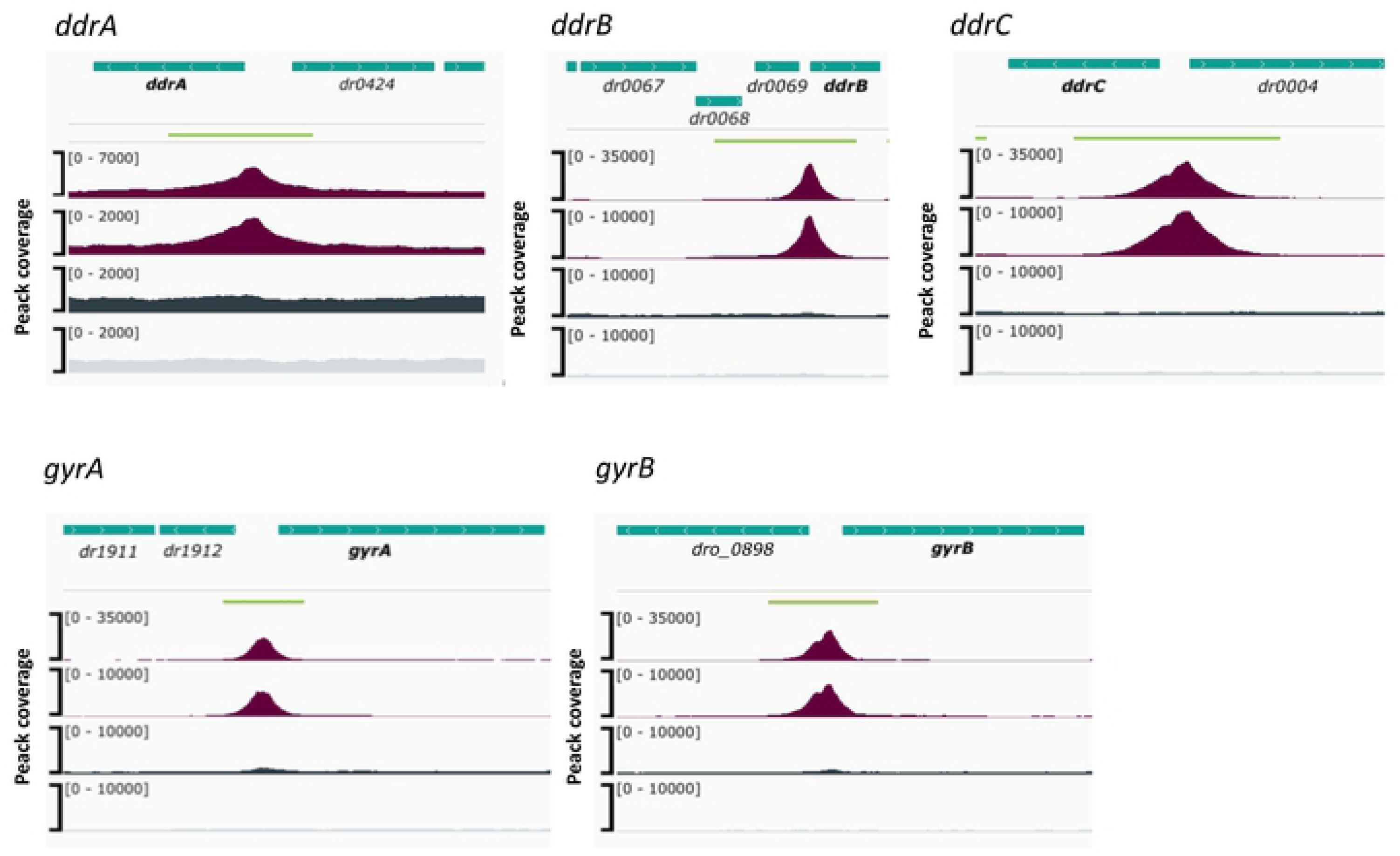
Visualization, through IGV, of the binding peaks obtained from genome analyze. Tag density profiles are illustrated for 2 IP (purple), the Input (dark grey) and the mock (light grey) for five known DdrO-regulated genes: *ddrA, ddrB, ddrC, pprA, gyrA and gyrB.* The green lines indicated the size of each peak identified by bPeaks. Genes are represented by green boxes, their location on either strand is indicated by > (strand +) and < (strand -).

To identify candidate binding sites of the DdrO protein, the nucleotide sequences of the ChIP peaks, between 151 and 1401 in length, were compared using MEME [38], to search for palindromic or non-palindromic motifs with an occurrence of one motif per sequence or any number of repetitions (Fig 1B). A total of 41 peak sequences, located in the promoter regions, contained a conserved DNA motif close to the RDRM sequence, with some loci containing two motifs (S2 Table). Interestingly, based on ChIP-seq data, the number of RDRM reported here is larger than that predicted by previous *in silico* analyses [19, 33]. No other conserved sequence pattern was found, except for the core promoters, either from the 41 ChIP peaks sequences or from the other sequences lacking an RDRM. However, a degenerate RDRM might not have been detected due to the threshold used for these bioinformatic analyses. Altogether, these results confirmed *in vivo* the role of the RDRM for DdrO binding to the *D. radiodurans* genome. Independently, we investigated whether an RDRM was found in the promoter regions of other genes encoded by *D. radiodurans*. We monitored, with FIMO, their presence in a set of sequences covering the regions located between -500 / +100 nucleotides from the start of translation of all the 3147 CDS encoded by *D. radiodurans*. A total of 222 putative RDRM-like sequences (Fig 1B) were found, including the 41 detected by MEME and 8 other potential sites, that were not detected by MEME, but 6 of which were located far from the start of coding sequences and outside the ChIP-seq peaks.

Based on ChIP-seq results, 89 genes, sometimes included in operons, may be regulated by DdrO, but many enriched peaks were located within the intergenic region of divergently transcribed genes. It is possible that only one of the two divergent genes may be under the control of DdrO.

### Transcriptome analysis of *D. radiodurans* in response to the depletion of DdrO

In parallel, to further characterize the RDR regulon in *D. radiodurans*, we compared transcriptome profiles of cells expressing, or not, DdrO. Since DdrO is essential for cell viability [17, 20], we used a conditional gene inactivation system [20, 39]. In this system, Δ*ddrO* cells expressed the DdrO protein under control of its own promoter at 30°C from a temperature-sensitive (*repU_T_*_s_) replication vector (Fig 3A) [20].

**Fig 3:**
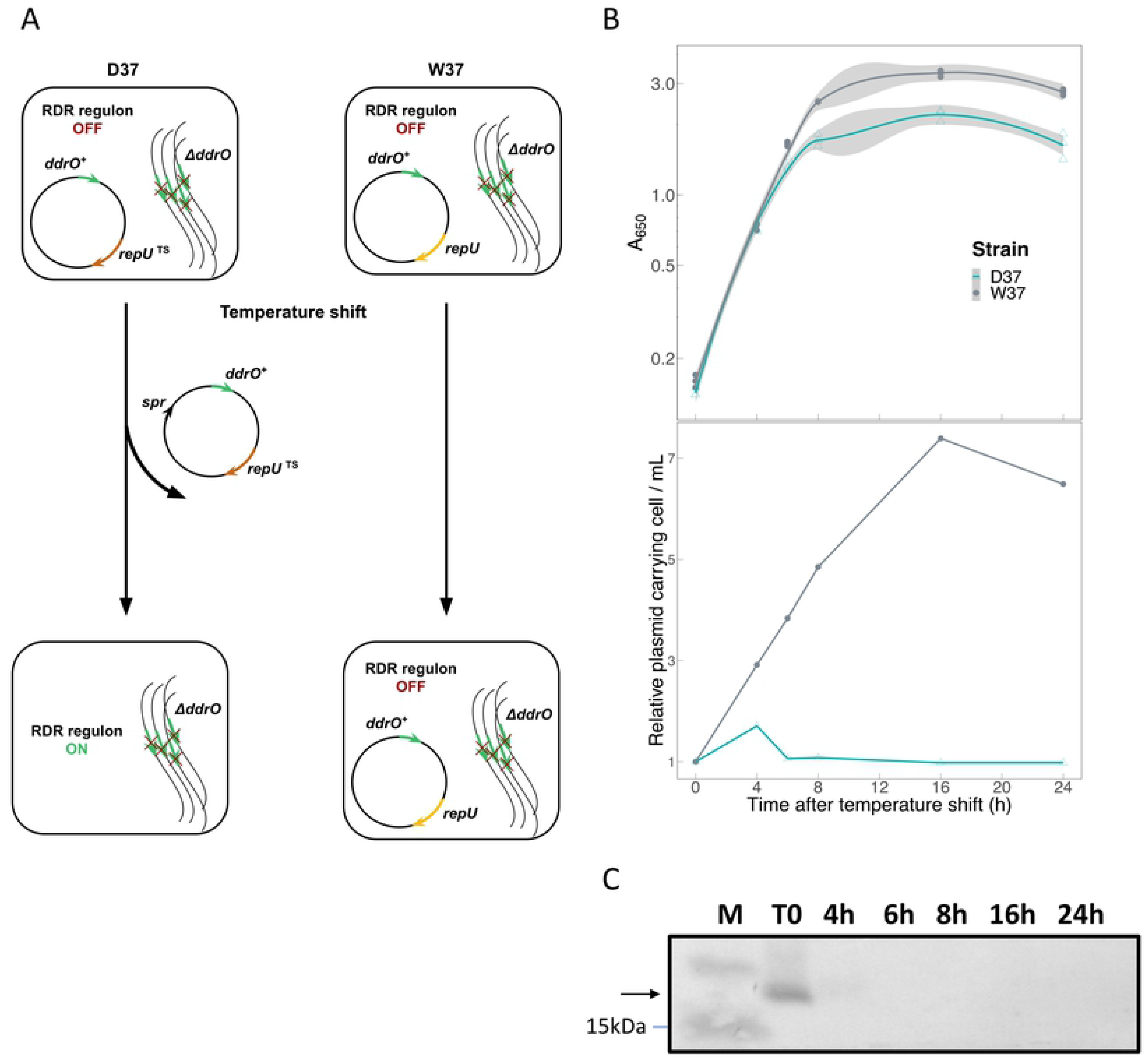
Loss of the *repU_TS_* vector in a chromosomal *ΔddrO* context. (A). Experimental design. Expression of RDR regulon genes was induced at 37°C when the thermosensitive replicative vector expressing DdrO could no longer to replicate. (B). Growth parameters of W37 and D37 strains, and relative stability of the *repU_T_*_s_ or the *repU*^+^ replication vectors expressing DdrO at 37°C during 24 h. The A_650_ values of the cultures were measured in 3 independent experiments. To calculate vector stability, samples were removed at the indicated times for plating at 30°C on media with or without spectinomycin. (C). Western-blot analysis of recombinant DdrO-FLAG protein depletion in D37 at 37°C. At each time point, aliquots of cells were removed and cell extracts (15 μg of proteins) were subjected to SDS-PAGE and analyzed by Western blot with anti-FLAG antibodies. The indicated times are relative to the initial temperature shift time point (0 h).

Shifting the culture to 37°C, a non-permissive temperature, resulted in an inability of the plasmid to replicate during successive cell divisions, leading to the depletion of DdrO, in contrast with a derivative of this expression vector, containing the wild type *repU*^+^ gene, that did not cause depletion of DdrO at 37°C [20, 39]. The Δ*ddrO*/*ddrO*^+^ (prepU*_Ts_*) and Δ*ddrO*/*ddrO*^+^ (*repU*+) strains grown at 37°C are denoted D37 and W37, respectively. Under our experimental conditions, the number of cells carrying the *repU^+^* vector is proportional to the increase of cell mass at 37°C without selective antibiotic (Fig 3B). In contrast, the number of cells carrying the *repU_TS_* vector stopped increasing and remained stable over 24 h (Fig 3B). The growth curves of both strains exhibited a comparable doubling time over 6 h. However, after this time lapse, the growth of the D37 strain also stopped, coinciding with the stress triggered by the depletion of DdrO (Fig 3B).

In a first attempt, we analyzed the effect of DdrO depletion on the expression of DdrD, DdrO PprA and RecA proteins belonging to the RDR regulon. For this purpose, we used derivatives of the strain Δ*ddrO* (p*repU_Ts_*::*ddrO*^+^) expressing DdrO-FLAG, PprA-HA, DdrD-HA or RecA-HA tagged proteins. Depletion of the DdrO repressor, in cells grown at the non-permissive temperature, resulted in the complete loss of DdrO after 4 h at 37°C (Fig 3C) and an increasing amount of PprA, DdrD, RecA proteins during the kinetics (S2 Fig).

In a second step the kinetics of gene expression changes induced by DdrO depletion were analyzed for both strains, from 3 independent cultures and at 6 time points (1 h, 4 h, 6 h, 8 h, 16 h, 24 h) (Fig 4A). RNA sequencing data was performed from 36 samples, corresponding to each sample to an average sequencing depth of 647-fold the genome sequence.

**Fig 4:**
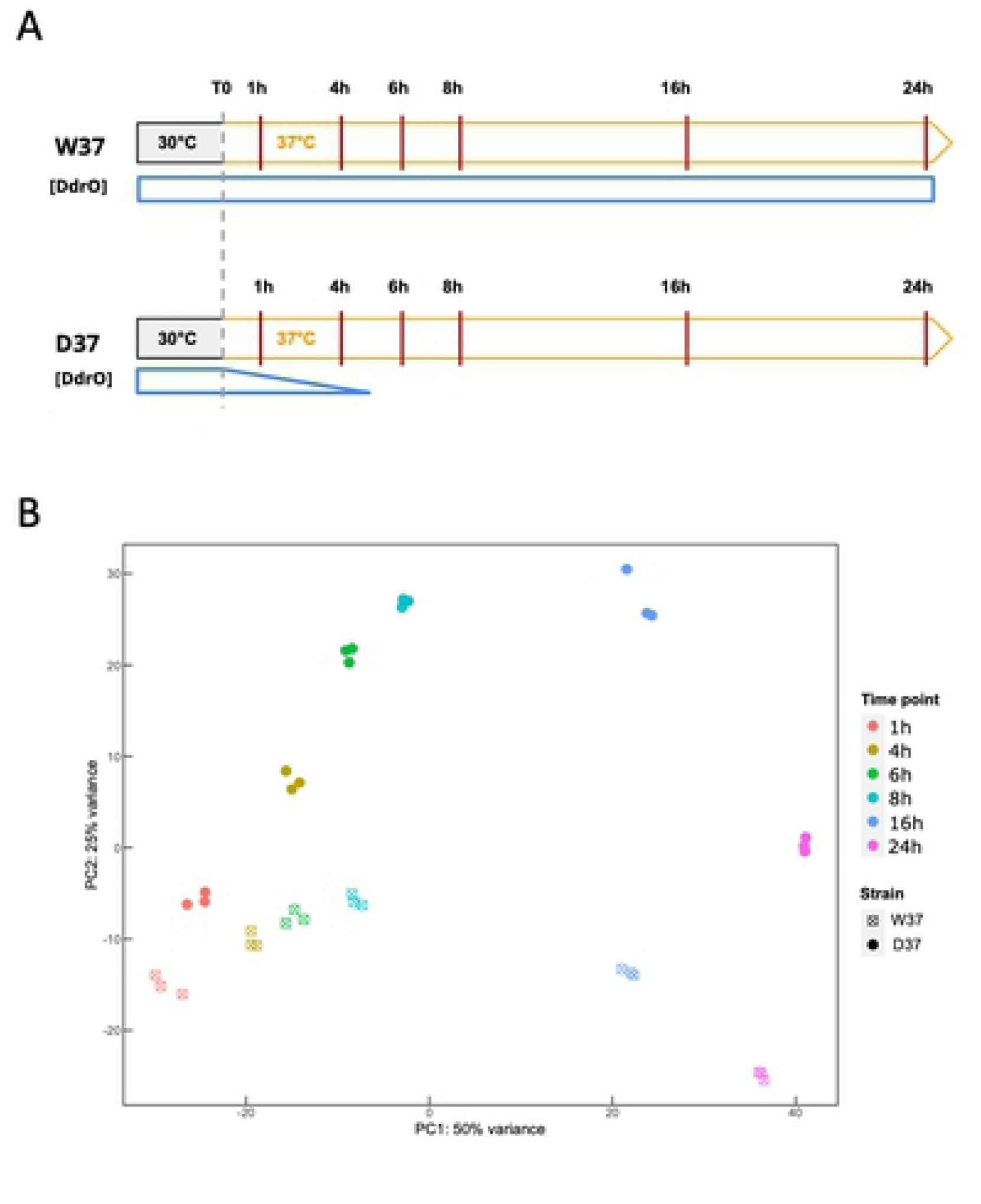
Transcriptomic analysis of DdrO depletion. (A). Schematic representation of the transcriptome time course. D37 and W37 cells were first cultivated at 30°C and exponentially growing cells were rapidly transferred to 37°C. The six time points examined are indicated by vertical lines. The amount of DdrO during the time course in both strains is indicated by blue plots. (B). Principal component analysis of all D37 and W37 samples after temperature shift. Each replicate was plotted as an individual data point. The indicated times are relative to the temperature shift time point (0 h).

A 2-fold change in expression threshold for the ratio in these experiments was applied, together with a *p-Value* < 0.01. Principal component analysis (PCA) confirmed that the transcriptome of the 3 biological replicates is clustered at each time point, showing the reproducibility of the experiments and the transcriptome patterns evolved as cells progressed through the time course of the experiment (Fig 4B). The data sets are separate, as well as the PC1 and the PC2 levels, which together explained approximately 75 % of the variance.

To compare the transcriptome, the 1 h time point was used as the reference, giving time for the genome to stabilize its expression after shifting the temperature. We first compared, for each strain, the deregulation of all genes along the time course. The results of the differential expression for all genes in W37 and D37 are presented in Fig 5 and S3 Table. After an incubation of 24 h at 37°C, numerous genes were deregulated, as 2129 unique genes i.e. 67.7% of all genes in W37 strain, and 2330 unique genes, i.e. 74 % in the D37 strain, were up- or down-regulated at least at one time point during the time course, showing that a cascade of cell regulation occurred into each strain over 24 h.

**Fig 5:**
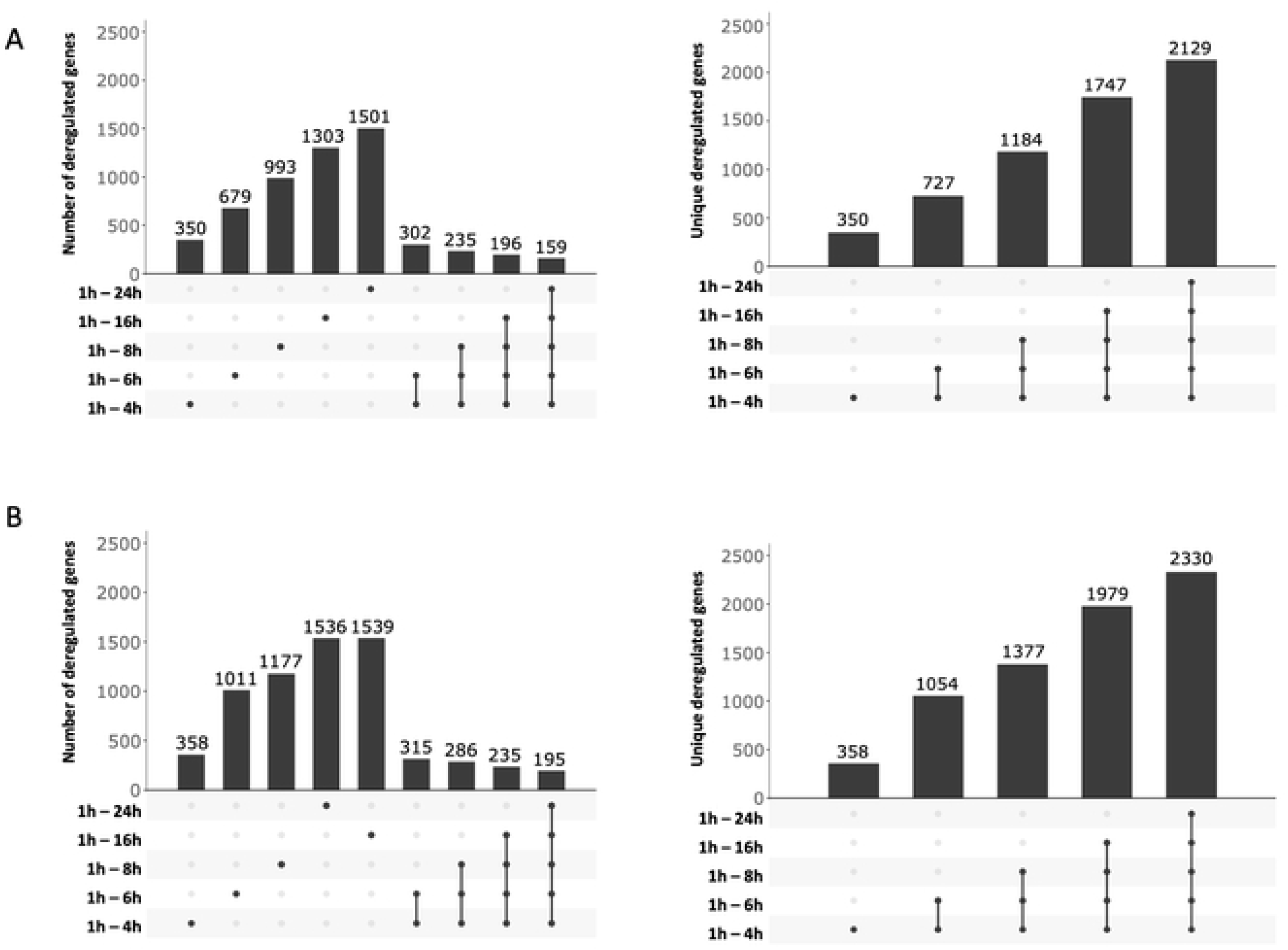
Cross match diagrams of deregulated genes from W37 (A) and D37 (B) strains after the temperature shift. The left panel shows the number of differentially expressed genes at each time point (S3 Table) and common genes among indicated time intervals (illustrated by solid black points, linked by black lines). The right panel shows the increasing number of unique genes that are differentially expressed at least at one time point after the temperature shift. Number of deregulated genes (|FC| ≥ 2, *pVal* ≤ 0.01).

Moreover, genes reported in one time-point were often found in the following time point. From the set of 350 and 358 regulated genes in W37 and D37 respectively during a time-point between 1 h and 4 h, 159 (45 %) in W37 and 219 (61%) in D37 remained regulated during all time points (S3 Table and Fig 5). After 6 h at 37°C, while most of the cells in the D37 strain had lost the thermosensitive plasmid, 679 and 1011 genes were deregulated in W37 and D37, respectively with 501 genes shared between them (S3 Table) and several upregulated genes found in W37 were downregulated in D37. These results showed that, rapidly after the temperature shift, the expression patterns of W37 and D37 changed differentially and that D37 cells likely implemented different cellular programs.

To confirm, at the transcriptome level the loss of the prepU*_Ts_* plasmid at 37°C, we investigated the expression profiles of *ddrO* and *spr* encoding resistance to spectinomycin. As shown in S4 Table, the *spr* gene was downregulated in D37 confirming the loss of the *repU_T_*_s_ plasmid in growing cells at 37°C (Fig 3) while the *ddrO* gene was upregulated in D37. When Δ*ddrO* / *ddrO*^+^ strains were constructed, only the CDS encoding the DdrO protein was deleted from the genome. Therefore, the upstream region, containing the promoter as well as the 5’UTR region of *ddrO,* is duplicated, one located on the chromosomal locus, the second on the plasmid. The RDRM is located in the *ddrO* gene 153 nucleotides upstream of the ATG in the vicinity of the predicted promoter. The 5’UTR reads density profiles of *ddrO* were very low in W37 strain, but were augmented in D37 strain as soon as cell lost the plasmid, supporting previous studies that DdrO regulates the expression of its own gene [17, 26] (S3 Fig).

Twenty genes of the RDR regulon were upregulated in the D37 strain, often from the beginning of the experimental temperature shift with increasing fold changes as cells progressed through the time course. The *dr2256* gene encoding a transketolase as well as the *ddrF* and *ssb* genes are also upregulated latter (6 h or 8 h, S5 Table) and the *uvrA* gene (*dr1771*) was upregulated only after 16 hours at 37°C. The other genes such as *uvrD* (*dr1775*) and *irrI* (*dr0171*), were not upregulated or only changed in a late stage of the experiment. On the other hand, *drA0151* encoding the first gene of the *hut* operon, was strongly downregulated in W37 strain and in D37 strain, but this gene was not reported as being under the control of the DdrO/IrrE proteins [24]. We also wondered if other *D. radiodurans* genes displayed a transcriptome pattern comparable to the RDR regulon genes. For this purpose, we selected genes as differentially expressed (DE) in W37 strain in less than or in two comparisons (DE ≤ 2), and in more than three comparisons in D37 strain (DE > 3) considering most of the profiles exhibited by the predicted RDR genes (see Methods, S2 Table). A total of 436 genes displayed similar transcriptome profiles (Fig 1C), reduced to 151 genes when an additional filter was applied to focus on genes which exhibited differential expression (up- or down-regulation) at only intermediate time points, *i.e.* 6 h, 8 h and 16 h. A total of 260, out of the 436 identified genes, were upregulated in the D37 strain (S6 Table), and 60% of these were distributed into four functional categories. A total of 31 genes encoding proteins involved in DNA metabolism and including most of the previously predicted RDR regulon, were found, but also the *recG* and *recO*, genes encoding a primosomal protein N’, as well as two DNA polymerase III subunits, *polA*, two putative helicases, *ligA*, *mutS* and *recN* (Fig 6). Interestingly, several genes encoding transcription factors involved in stress responses as LexA1, LexA2 or the Phage Shock Protein A PspA [40] were also upregulated in D37 (Fig 6). Therefore, several regulatory networks were likely triggered in response to induction of the RDR regulon.

**Fig 6:**
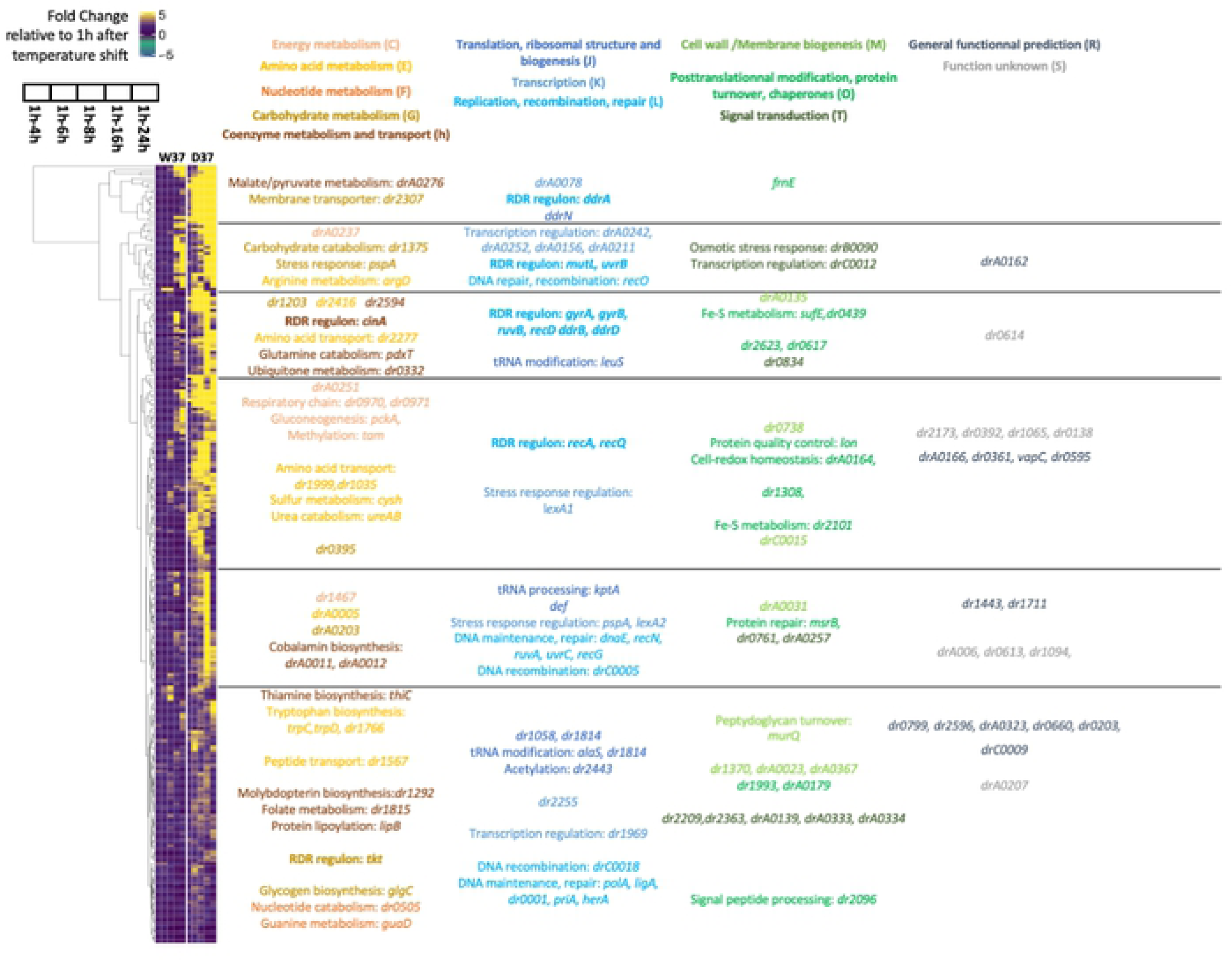
Hierarchical clustering of genes specifically upregulated in response to DdrO depletion. The 260 genes differentially expressed between time segments 1 h-6 h and 1 h-16 h in the W37 strain in less than two comparisons (DE ≤ 2) and in more than three comparisons in D37 strain (DE > 3) were hierarchically clustered according to their temporal expression. Only several genes representing some COG categories are shown.

In addition, 15 genes encoding putative proteases and peptidases or regulators of protease activity, 15 genes coding for ABC transporters, permeases and efflux components and > 100 genes coding for uncharacterized proteins or of unknown function were also deregulated with similar expression patterns (Fig 6).

DdrO was described as a transcriptional repressor [17, 20], but one cannot exclude that this regulator is also able to enhance several genes expression under normal growth conditions. Therefore, we also investigated the presence of downregulated genes during DdrO depletion. Using the same settings, 176 downregulated genes were found (S4 Fig and S7 Table), widely distributed in the different COG categories, with a transcriptomic repression mainly beginning at 6 h when D37 cells are depleted in DdrO.

Our RNA seq analysis exhibited 436 deregulated genes, but the overall up- or down-expression of all these genes may be a consequence of a cascade of regulation occurring when cells lost the *ddrO* gene.

### Integration of the data: the DdrO map in *D. radiodurans*

To map the DdrO regulon, we integrated the results obtained in ChIP-Seq, RNA-seq and motif research. When candidates from these three experiments were compared, 37 genes met all three criteria, and 48 other genes met two criteria (Fig 1D and Fig 7). Based on these results, the DdrO regulon comprises 16 previously predicted RDR genes, mainly involved in DNA repair pathways. Moreover, 3 other genes involved in DNA metabolism (*recG, helD* and DNA ligase *ligA)*, 4 genes associated with different metabolic pathways, 5 genes involved in translation, as well as 7 new genes encoding proteins of unknown function (Fig 7, Table 2 and S8 Table) are also under the control of DdrO.

**Fig 7:**
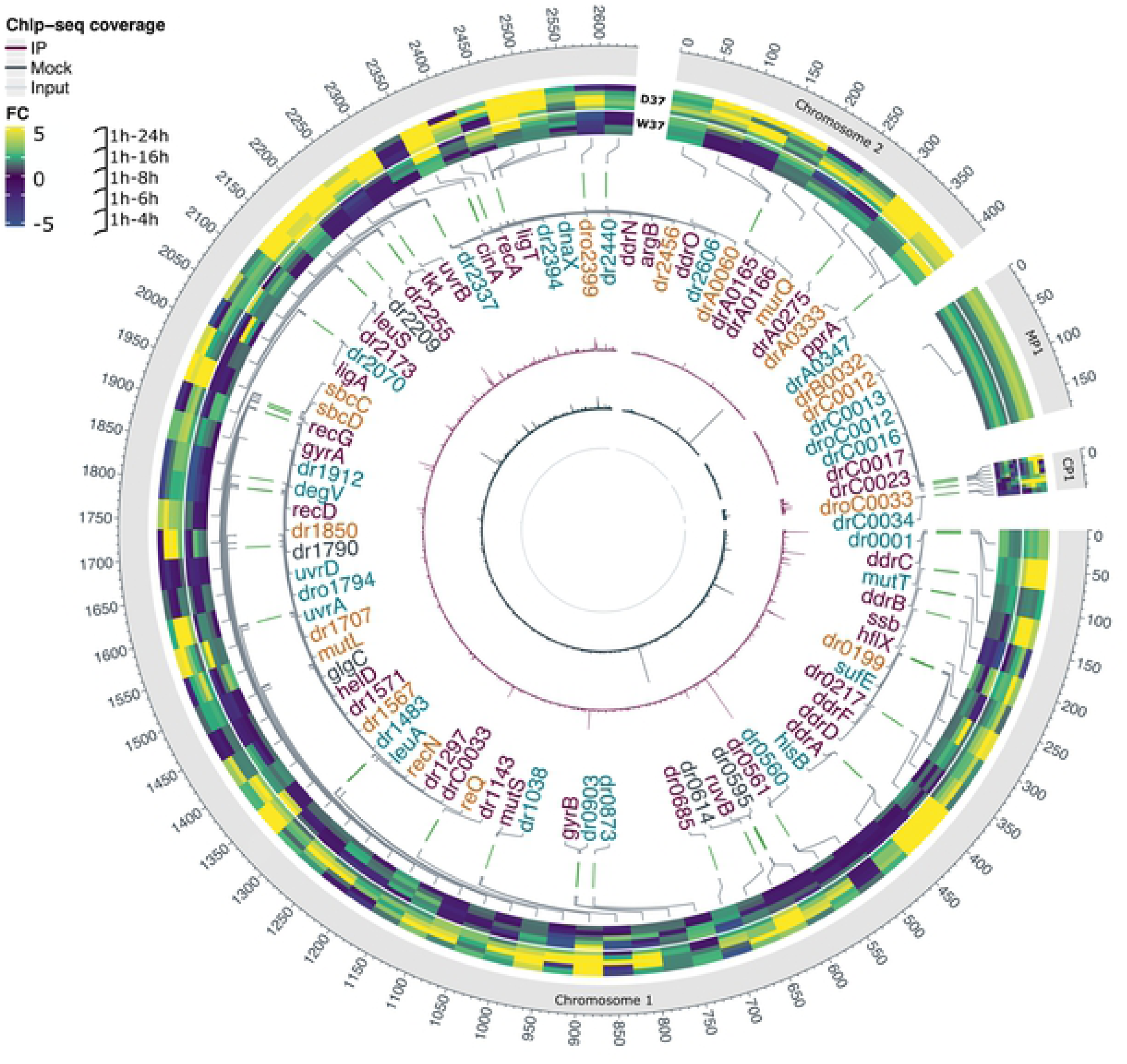
Representation of identified DdrO target genes across the *D. radiodurans* genome. Each genome replicon is represented by an outer circle. Heatmaps represent Fold Change values for the 85 genes, sometimes in operons, matching with at least 2 of the selected criteria (see Fig 1). The genomic positions of deregulated genes are drawn in grey connections. The DdrO bound sites associated with identified candidate genes included in the DdrO regulon are illustrated by green vertical lines. High confidence DdrO targets genes matching with 3 criteria are indicated in purple. The other genes matching with two criteria are labeled in dark gray (ChIP-seq and Transcriptomics) in blue (Transcriptomics and RDRM) or in orange (ChIP-seq and RDRM). The purple circle shows the mean coverage from the three IP replicates. The dark grey circle shows Input tag density profiles and the light grey shows the mock tag density profiles.

**Table 2:**
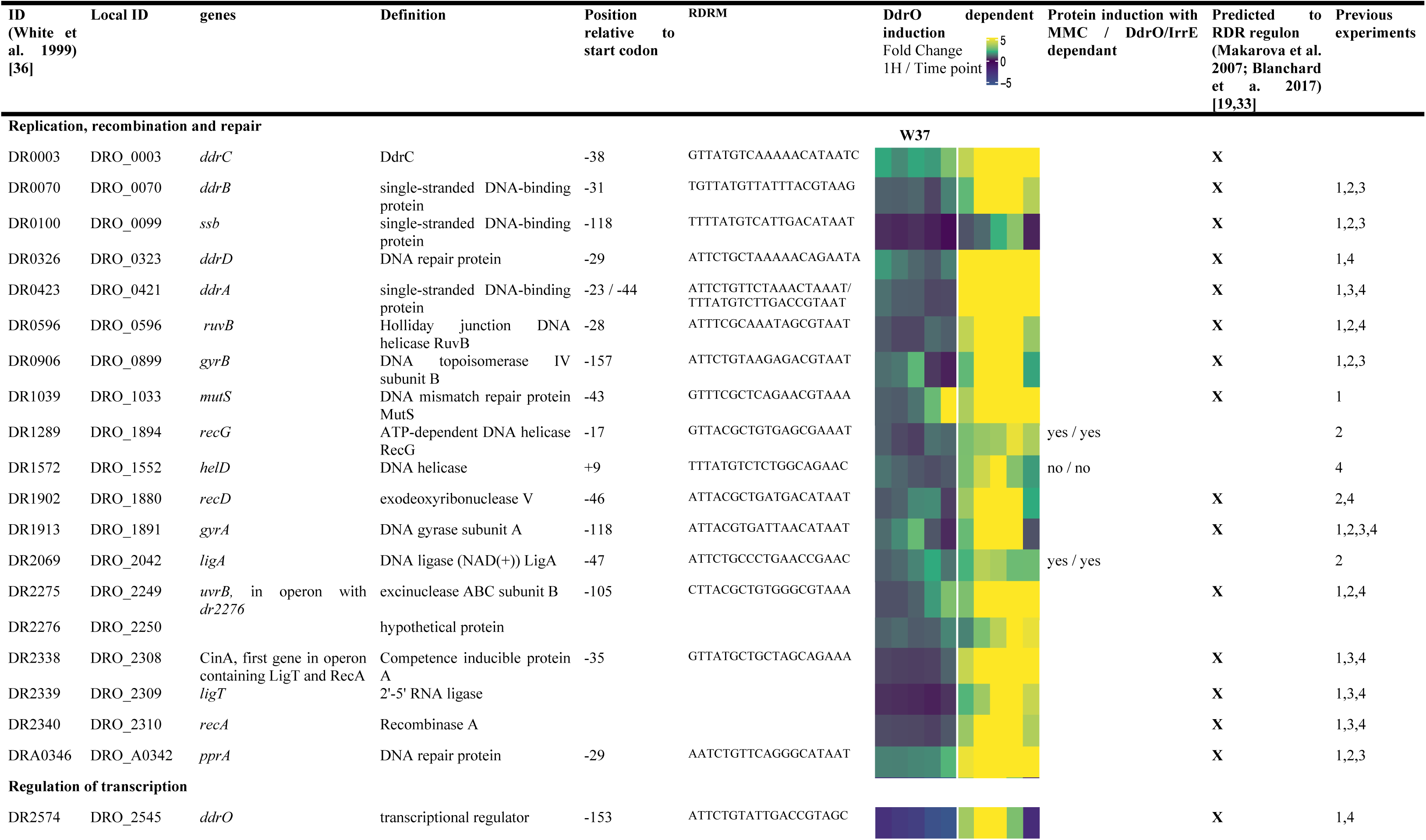
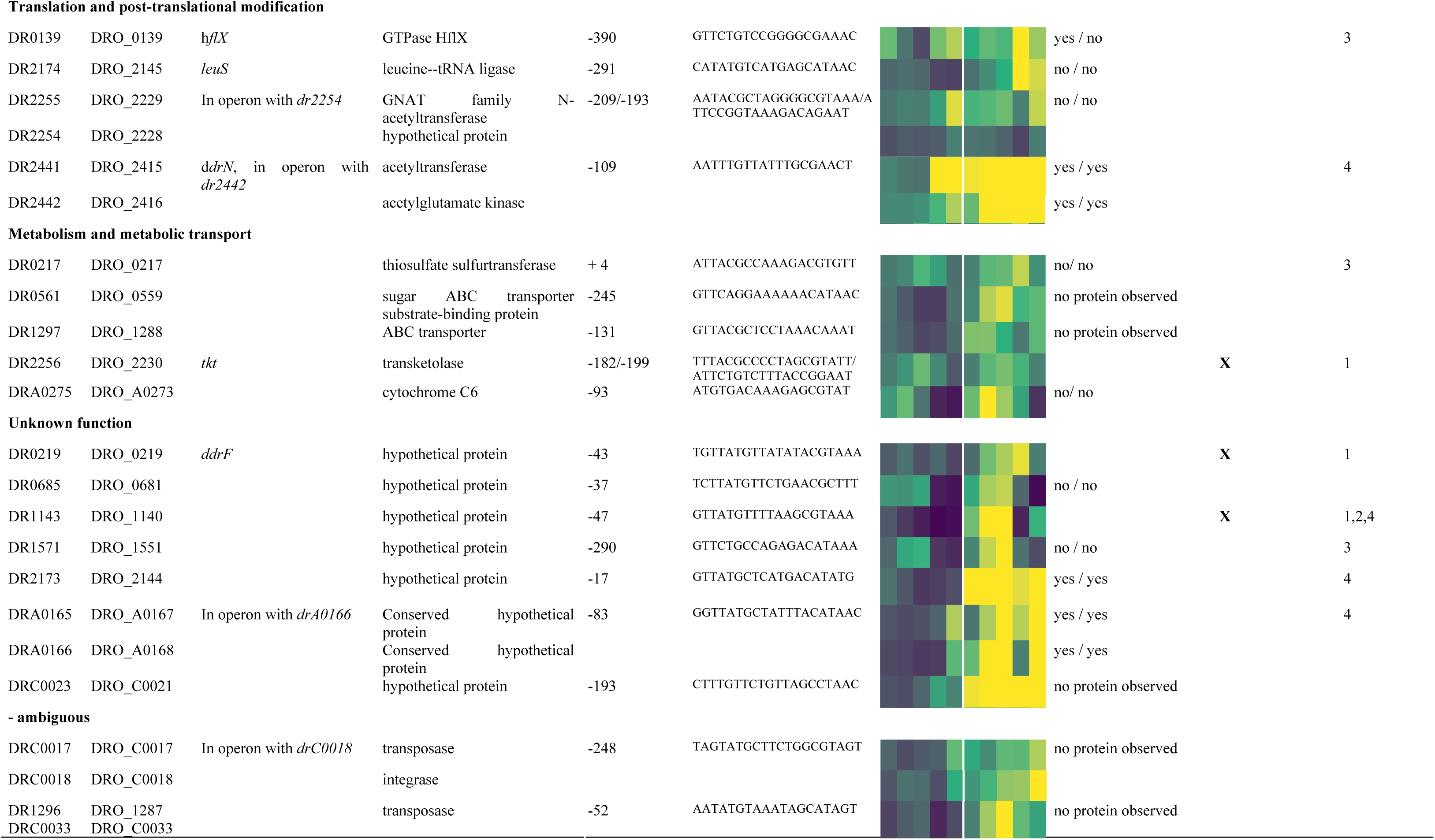
List of gene belonging to the DdrO regulon in D. radiodurans matching all criteria (transcriptomic, ChIP-seq and RDRM). RDR genes previously predicted by bioinformatic analyses and by EMSA assays (1) [26, 33] or shown to be regulated after exposure to radiation by transcriptomic (2) [12], proteomic studies (3) [41] or in a *ΔirrE* mutant (4) [24], respectively.

Surprisingly, two genes, *dr0017* and *d1287,* encoding transposases also matched all the criteria. One copy of *drC0017* and two copies of the second IS (*dr1287* and *drC0033*) are present in the genome. According to the Chandler classification (http://www-is.biotoul.fr/) *dr1287* encodes the ISDra5 and *drC0017* is part of the transposable element TnDra1. Since *drC0033* and *dr1287* CDS exhibited 100% sequence identity, we were not able to determine, by RNA-seq, whether one or the two genes displayed an upregulation during the time course. Moreover, only a small peak was observed for *drC0033* but was below the threshold we established for this study.

In addition, 48 genes matched only two criteria. Among them, 6 genes were upregulated in D37 and a ChIP peak was located in each promoter, but their sequences did not exhibit any RDRM or other conserved motif in their core promoter region (S9 Table). However, we cannot exclude the possibility that the DdrO protein bound to a degenerate RDRM sequence, that were not reported because of the criteria used here for *in silico* analyses. Among them*, dr2606* encodes a predicted primosomal protein N’ and *dr1790* encodes for the yellow protein, belonging to the ancient yellow/major royal jelly (MRJ) protein family. The deletion *of dr1790* in *D. radiodurans* increased its membrane permeability and decreased cell growth rate and survival upon exposure to hydrogen peroxide and radiation [42].

Twenty six other genes were only associated with a DdrO peak and also with an RDRM in their promotor region (Fig 7 and S10 Table). From this set of 26 genes, 13 are divergently transcribed from genes that matched all criteria. It is thus likely that only one of the two genes that share the same intergenic region were regulated by DdrO. Seven out of the 13 other genes, such as *dr0001* encoding DNA polymerase III subunit beta, were described as upregulated upon exposure to gamma rays [12] but was not reported as differentially expressed in a *ΔirrE* mutant [24]. The *uvrA* and *uvrD* genes were also reported as being upregulated in the first 1.5 h when cells are exposed to large doses of gamma-rays [12] and were differentially expressed very early in a *<u>Δ</u>irrE* mutant. These results suggest that the expression of these genes, including *uvrA* and *uvrD,* may be augmented very early during the time course (< 1 h) or are under the control of DdrO and other regulatory elements, and the simple depletion of DdrO did not modify their expression under our experimental procedures.

Finally, 16 genes were only found to be upregulated in D37 and not in W37, and an RDRM was detected near their respective core promoter but no ChIP-Seq peak was reported from the ChIP-seq analysis (S11 Table). However, careful inspection of all the peaks with the IGV program showed that a small peak, that fell below the threshold used to analyze ChIP-seq data, was observed in the promoters of 5 genes (*sbcD*, *recQ, drC0012* encoding a putative transcriptional regulator*, drC0033* (transposase) and *dr1707* encoding DNA polymerase I). A highly condensed structure of the *D. radiodurans* chromatin may have locally impaired or decreased the efficiency of the ChIP experiments, and thus, these genes might also belong to the DdrO regulon.

The location of the RDRM was also analyzed in the promoter region of all genes matching two or three of our criteria. The RDRM sequences were mainly located in the vicinity of the predicted position of *E. coli*-like −35 and −10 promoter consensus sequences (S8-S10-S11 Tables). These results are consistent with previous reports showing that RDRM in *D. deserti* was found both upstream (−50 bp) as well as downstream (+20 bp), of transcriptional start sites (TSS) potentially overlaping with RNA polymerase binding site [17].

In parallel, the amount of protein encoded by the newly identified genes was analyzed in wild-type cells and in a *ΔirrE* mutant after exposure to mitomycin C (MMC). For this purpose, we monitored, by Western-blot analysis, the expression of C-terminal -tagged recombinant proteins. The cellular levels of 5 recombinant proteins (RecG, LigA, DdrN, DRA0166 and DR2173) increased in wild-type cells in response to MMC but remained constant in an *ΔirrE* mutant, thus corroborating our data (Fig 8). We were not able to detect *dr1297*, *dr0561* tagged proteins, but these genes encode a predicted ABC transporter and a sugar ABC transporter respectively, containing predicted transmembrane regions or a predicted periplasmic peptide signal that could lead to a low solubility of these proteins. In agreement with our gene expression data, previous transcriptome studies have shown that *dr0561* was indeed not upregulated in a *ΔirrE* strain when compared to a wild-type strain upon exposure to γ-ray irradiation [24] lending support to the idea that this predicted transporter is regulated in a DdrO/IrrE manner.

**Fig 8:**
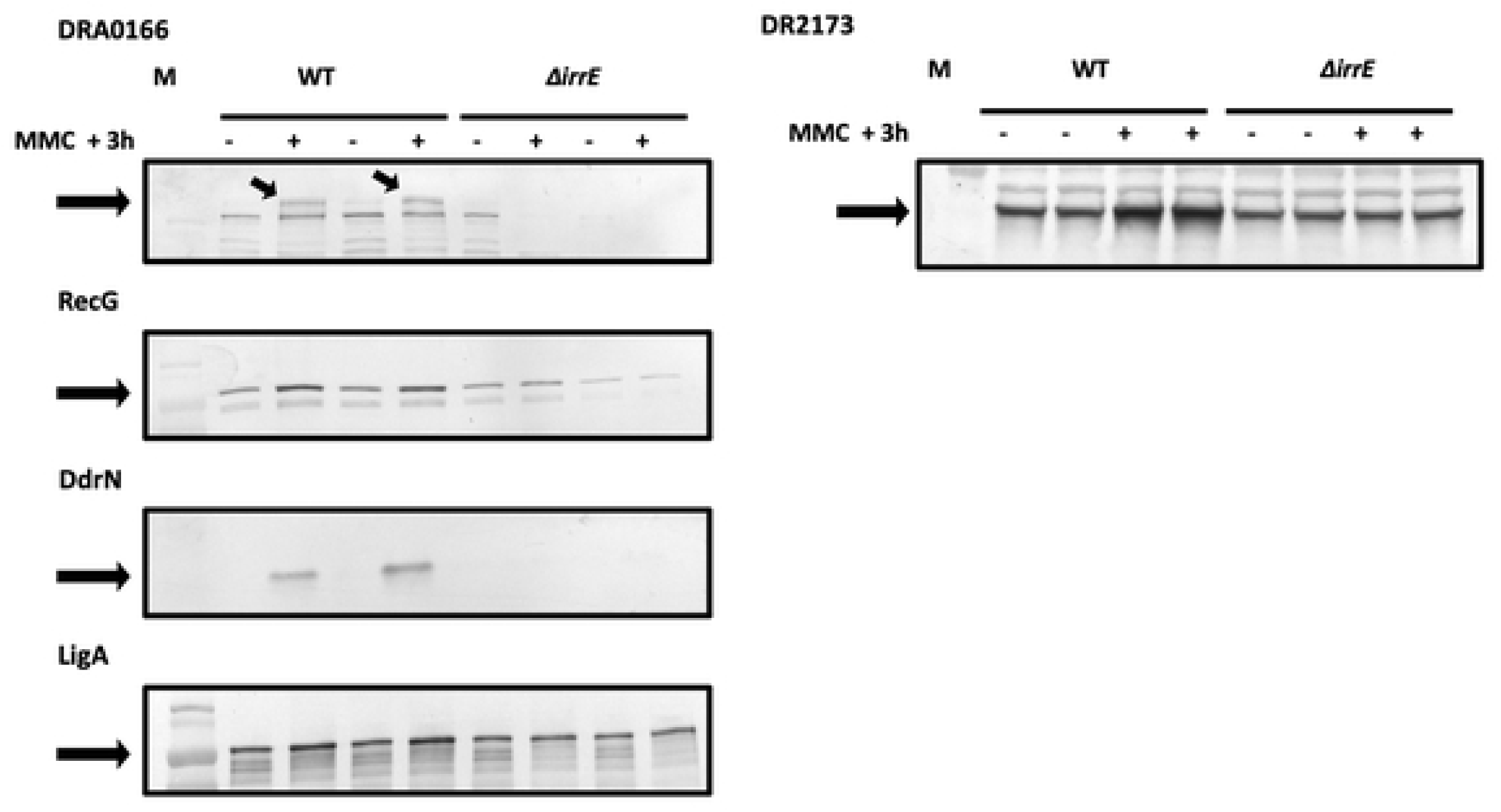
Expression of several new DdrO targets genes induced after exposure to MMC in an IrrE and DdrO-dependent manner. *ΔirrE* or wt cells expressing recombinant DRA0166-V5, RecG-V5, DdrN-V5, LigA-V5 and DR2173-V5 proteins, respectively were incubated (+) or not (-) with MMC (1 µg/ ml) at 30 °C for 3 h. Cell extracts were subjected to SDS-PAGE and analyzed by Western blotting with anti-V5 antibodies. For DRA0165 and DR2173, 15 μg of proteins were loaded on each well, for RecG and DdrN, 10 μg of proteins, and for LigA, 5 μg.

Among the set of genes predicted *in silico* by Makarova et al [33], the *frnE* and *rsr* genes were upregulated over time course experiment in D37 but no DdrO-peak was found in their respective promoter regions. We analyzed the expression of the FrnE and Rsr proteins and also of DdrR (DR0053) predicted as belonging to the RDR regulon in *D. deserti* [19] although *ddrR* was not upregulated during our time course experiment and no peak was observed in its promoter region. The cellular levels of three recombinant proteins increased in response to MMC in wild type cells, and also in a *ΔirrE* mutant (S5 Fig). These results strongly suggest that DdrR, FrnE and Rsr proteins are induced by genotoxic stress but not in a DdrO/IrrE dependent manner.

Except for HflX, which was found to be induced in wild-type and in *ΔirrE* cells after exposure to MMC, no change in protein quantity was observed for DRA0275, DR0217, DR0685, DR1571, DR1572, DR2255 and DR2174 after exposure to MMC (S5 Fig). However, as already reported in *D. radiodurans,* the upregulation of several genes at the transcriptomic level was not always observed at proteomic level [41]. This could be due to their abundance in the cell, their very transient expression or their instability. A mechanism of translational regulation might also occur after transcription of these genes. Alternatively, it is possible that the genotoxic conditions after MMC exposure did not exhibit appropriate deleterious effects in cell to trigger induction of these proteins.

## Discussion

An RDR regulon was proposed several years ago based on the presence of the RDRM sequence, a common 17 bp palindromic sequence, located in the promoter region of the most highly ionizing radiation and desiccation up-regulated genes in *D. radiodurans* and *D. geothermalis* [33]. To date, identification of putative DdrO target genes in *D. radiodurans* have been mostly proposed by a combination of bioinformatic analyses based on microarray genes expression data [33] and a validation, *in vitro*, by Electrophoretic Mobility Shift Assay (EMSA] experiments [26]. Here, we combined two large scale approaches to identify DdrO targets *in vivo* with a reliable accuracy. Analysis among transcriptome data, identification of enriched DdrO binding sites and the presence of an RDRM in the promoter region of *D. radiodurans* genes allowed us to identify at least 35 DdrO target genes matching all criteria (Table 2) and to propose other genes which may also be regulated by this transcription factor (S9-S10-S11 Tables).

Up to 70 % of the identified target genes were previously predicted to be part of the RDR regulon [19,26,33]. The *ddrF* gene (*dr0219*), absent of from the genome annotation published by Hua and Hua [35], is identified here as belonging to the RDR regulon, as initially described [33]. In addition, we highlighted 18 new DdrO target genes including genes involved in DNA maintenance such as *dr1289, dr1572 and dr2069* encoding the RecG helicase, HelD superfamily I helicase IV and the replicative DNA ligase LigA, respectively. In agreement with these data, transcriptome analysis of cells recovering from exposure to ionizing radiation showed no upregulation of expression of these three genes in a *ΔirrE* mutant compared to the wild-type strain [24]. In *E. coli*, RecG plays an important role in DNA repair, recombination and replication [43] while HelD, from *Bacillus subtilis* or *Mycobacterium smegmatis,* is associated with transcriptional pathways [44, 45]. In *D. radiodurans,* cells devoid of RecG exhibit a delay in growth, in double strand break (DSB) repair kinetics and a decrease in resistance to γ-irradiation and H_2_O_2_ [46, 47] whereas the *Δdr1572* mutant exhibited a greater sensitivity to H_2_O_2_ but no change in resistance to ionizing radiation and to MMC when compared to the wild-type strain [48].

The identification of *ligA* as a target gene is interesting, since DNA ligases are implicated in DNA repair and are essential in other fundamental processes within the cell [49]. Ligase activity is crucial during DNA recombination and replication explaining the constitutive expression of DNA ligase during all phases of the cell cycle [49]. Therefore, DdrO binding on the *ligA* promoter region should not completely repress gene expression, to ensure so as a minimum level of DNA ligase activity. However, in response to elevated amounts of DNA damage and particularly to DSB, the basal level of LigA may not be sufficient for accurate Single Strand Annealing (SSA) [8] and Extended Synthesis-Dependent Strand Annealing (ESDSA) mechanisms, as well as for homologous recombination [5].

We identified new DdrO target genes, *ddrN* and *dr2255* encoding putative GNAT family acetyltransferases that may be involved in post-translational modification (PTMs) pathways. The *ΔirrE* cells recovering from exposure to ionizing radiation exhibited no upregulation of *ddrN* expression when compared to a wild-type strain [24]. PTMs in bacteria play crucial roles in various cellular pathways, including after metabolic shifts and stress adaptation [50]. Acetylation is known to modify a variety of substrates involved in RNA metabolism, enzymatic activity or DNA related mechanisms[51]. In *E. coli*, acetylation of the chromosomal replication initiation protein DnaA leads to an arrest of DNA replication [52]. Moreover, acetylation of histone-like nucleoid protein HU in *M. tuberculosis* alters the *in vitro* DNA-binding capacity of HU as well as DNA structure, which may affect gene transcription and other protein - DNA interactions [53]. *D. radiodurans* HU protein has been reported as a major actor of nucleoid compaction [39, 54] and it may also be acetylated. Further analyses are required to understand the impact of acetylation activity in response to DNA damage in *D. radiodurans*.

Surprisingly, two genes, *drC0017* and *dr1287,* encoding transposases belonging to the Tn3 family and to the ISDra5 family, respectively, were identified as belonging to the DdrO regulon. Transposons are major actors of genome remodeling and play an important role to create diversity and to facilitate adaptation of the host to extreme environmental conditions. Insertion sequences are abundant in *D. radiodurans* and IS transposition is a major event in spontaneous, as well as induced, mutagenesis [55]. It has been previously shown that ISs belonging to different families (ISDra2, ISDra3, ISDra4, ISDra5, IS2621) were transpositionally active in this organism under normal growth conditions and transposition was enhanced in cells recovering from DNA damage. Transposable element expression and movement are generally tightly regulated and different mechanisms control their gene expression. In *E. coli*, LexA protein represses expression of the Tn5 transposase gene [56]. Further studies are required to better understand how DdrO contributes to the regulation of transposition events of these two ISs.

We also identified several genes encoding proteins of unknown function, such as *dr2173* and the *drA0165-drA0166* operon which were strongly upregulated in response to a depletion of DdrO. These genes were not upregulated in irradiated *ΔirrE* cells [24]. Domains of unknow function (DUF) found in DR2173 and DRA0166 are widely conserved in bacteria. A DUF4132, within DRA0166 may be involved in the molybdopterin biosynthesis. DR2173 also shares an N-terminal WGR domain with the MolR protein, which may be involved in regulation of molybdate biosynthesis in *E.coli* [57] and was described as belonging to the LexA regulon [58]. However, molybdate-metabolism associated genes, such as *D. radiodurans moeA* or *moeB,* were not differentially expressed in D37 compared to W37. The role of these strongly upregulated unknown genes in response of DNA damage also remains to be discovered.

DdrO-structure and biochemicals data suggested that in response to DNA damaging condition, upregulation of the expression of genes of the RDR regulon is dependent on a dynamic balance between DdrO dimers bound to DNA and the IrrE-cleavable DdrO monomer forms [25, 27]. Thus, the cleavage of DdrO monomers by IrrE would reduce the amount of DdrO dimers able to bind to DNA, leading to the induction of expression of genes controlled by this regulator. Here, the transcriptome data showed that identified RDR regulon genes were not all upregulated at the same time during the DdrO depletion. The expression of some genes such as *ddrC, ddrD* and *pprA* were strongly upregulated at early times whereas *ssb, uvrD* or *ddrF* showed a late upregulation suggesting that DdrO bound with more or less affinity to DNA according to the divergent RDRM sequences that may diverge. Therefore, after exposure of ionizing radiation, some genes would be upregulated earlier than others during cell recovery. It has been shown that, following extended DdrO-depletion, *D. radiodurans* cells were engaged in an apoptotic-like cell death (ALD) pathway leading to morphological alterations such as larger cells, membrane blebbing and DNA fragmentation [20]. In *Caulobacter crescentus*, the BapE endonuclease was reported to be involved in DNA fragmentation upon severe and extensive DNA damage [59]. BapE induction is triggered only in the case of prolonged LexA self-cleavage and was not described as part of early induced SOS response genes [59]. Instead, our results did not allow us to identify new genes belonging to the RDR regulon and differentially expressed at late times suggesting that ALD would be triggered by long-lasting induction of one or more genes from the RDR regulon, or by a cascade of regulation events following depletion of DdrO.

Despite the identification of many promoter regions containing a putative RDRM sequence in the *D. radiodurans* genome, we showed that only a small proportion of these are indeed bound by DdrO. Based on the presence of the RDRM sequence and induction of expression in response to ionizing radiation, *rsr, frnE, irrI, ddrR* and the *hutU* operon were previously described as part of the RDR regulon in *D. radiodurans* [19, 33]. In agreement with Wang et al [26], we showed that expression of the *hutU* operon, *rsR*, *frnE* and *irrI* is not under the control of DdrO, but also *ddrR*. However, we showed that the amount of RsR, FrnE and DdrR proteins is indeed induced in response to exposure to MMC (S5 Fig) in concordance with gene expression data [12,13,19]. Thus, identification of the RDRM sequence is not sufficient to enable DdrO-binding. Many factors, such as DNA structure or binding competition between multiple transcription factors can affect accessibility to a genome region for DdrO [60]. If our data showed that DdrO appears to bind exclusively the RDRM sequence, then some RDR regulon genes such as *ddrA, gyrA* or *dr1572* were also described as under the control of another major regulator, DdrI [61] highlighting the regulatory cross-talk in the *D. radiodurans* DNA damage response.

The *D. radiodurans* genome encodes more than one hundred predicted transcriptional regulators, but only a few studies have been performed to decipher genes that they may regulate. The RDR regulon of the radioresistant bacterium *D. radiodurans* characterized here by integrative genomic analyses, paves the way for further studies to better depict the regulatory networks underlying mechanisms that contribute to the extreme radiation tolerance of this fascinating bacterium.

## Material and Methods

### Bacterial strains, plasmids, oligonucleotides, media

Bacterial strains and plasmids are listed in S12 Table. The *E. coli* strain DH5α was used as the general cloning host, and strain SCS110 was used to propagate plasmids prior to transformation of *D. radiodurans* [62]. All *D. radiodurans* strains were derivatives of the wild-type strain R1 ATCC 13939. Transformation of *D. radiodurans* with PCR products, genomic or plasmid DNA was performed as previously described [6]. Strains expressing V5-tagged proteins were constructed by the tripartite ligation method as previously described [63]. The genetic structure and purity of the mutants were verified by PCR and sequencing. The sequences of oligonucleotides used for strain construction and diagnostic PCR are available upon request. Chromosomal DNA of *D. radiodurans* was extracted using the NucleoSpin DNA Microbial Mini kit (Macherey-Nagel). *D. radiodurans* genomic DNA used to sequence the genome with Nanopore technologies was prepared by a lysis procedure involving a pretreatment of the cells with saturated-butanol in EDTA [64]. PCR amplification of DNA fragments, using plasmid or genomic DNA as a template, was performed using Phusion DNA polymerase (Thermo Scientific).

*D. radiodurans* strains and derivatives were grown at 30°C or 37°C in TGY2X (1% tryptone, 0.2% dextrose, 0.6% yeast extract), or plated on TGY1X containing 1,5% agar, and *E. coli* strains were grown at 37°C in Lysogeny Broth. When necessary, media were supplemented with the appropriate antibiotics used at the following final concentrations: kanamycin, 6 μg/mL; chloramphenicol, 3,5 μg/mL; hygromycin 100 μg/mL; spectinomycin, 90 μg/mL for *D. radiodurans*; chloramphenicol, 25 μg/mL; spectinomycin 50 μg/mL for *E. coli*.

### *D. radiodurans* R1 sequencing, assembly and annotation

Purified *D. radiodurans* genomic DNA from strain R1 ATCC13939 (laboratory stock) was sequenced using Illumina NextSeq v. NS500446 (High-throughput sequencing facility of I2BC, Gif sur Yvette, France), yielding 10.7×10^6^ 75 nt paired-end reads. This dataset was subsequently assembled with SPAdes [v3.13.1] [65], with the ‘--carefull’ option set, and produced a total of 3255298 nt. in 136 contigs (N50: 149391).

In parallel, *D. radiodurans* genomic DNA was also sequenced with Oxford Nanopore Technologies GridION [v. GXB02022 – 19.12.6] (High-throughput sequencing facility of the I2BC, Gif sur Yvette, France), yielding a total 10.87 x 10^9^ nucleotites in 946434 long reads (median size: 6857 nt). This reads pool was further filtered with filtlong [v. 0.2.0] (https://github.com/rrwick/Filtlong), retaining only reads longer than 2000 nt, aligned to reference illumina reads (see above) and totaling ∼5×10^9^ nt. This reads dataset was used as input to Canu [v1.8] [66] for reads correction and assembly with default parameters, producing 4 low quality contigs, totaling 3578820 nt.

These latter sequences were improved by mapping SPAdes-produced high quality contigs on them with BWA mem [v. 0.7.9a-r786] [67]. SPAdes contigs alignments on nanopore reference sequences were extracted with samtools mpileup [v. 1.8] [68] and variants were called with bcftools call [v. 1.10.2] [69, 70] using prior probability set to 1 (-P 0.99). Lastly, terminal repeats were manually trimmed when present, and the sequence origin was adjusted to correspond to the sequence of White et al. [36] (AE000513.1, AE001825.1, AE001827.1 and AE001826.1) for each chromosomal element and plasmid.

CDS prediction was performed on the final assembled sequences using Prodigal [v. 2.6.3] using single mode, translation table 11 [71] as settings. Amino acid sequences of predicted genes were searched for similarity with BLASTP [72] to sequences from two other available complete sequences of *D. radiodurans* R1 strain (GCA_000008565.1 and GCA_001638825.1). Structural RNAs were mapped on the genomic sequences with the same reference genomes using the BLASTN tool.

### Time course experiment

#### DdrO depletion

*ΔddrO* strains complemented by *ddrO* expressed, under its own promoter, from a plasmid with wild type or thermosensitive (prepU_ts_) replication, were grown at a permissive temperature (30°C) with spectinomycin and chloramphenicol. Cells were diluted in fresh medium with antibiotics and grown at permissive temperature (30°C) and cells in exponential growth (A_650nm_ ∼ 0.5) were harvested by centrifugation, washed two times with TGY2x and reused to A_650nm_ = 0.1 in fresh medium without antibiotics. Then, temperature was shifted at 37°C (non-permissive temperature for the thermosensitive replication plasmid). At 1 h, 4 h, 6 h, 8 h, 16 h and 24 h, aliquots of 20 mL were removed for fluorescence microscopy and transcriptome analysis or Western-blot analysis.

#### RNA extraction, cDNA library construction and sequencing

For each aliquot, total RNA was isolated using the Fast RNA Pro Blue Kit (MP Biomedicals) and the FastPrep-24 instrument, according to the manufacturer’s protocols. Extracted RNA was rigorously treated with TURBO DNA-*free* (Invitrogen), according to the manufacturer’s instructions and the absence of DNA genomic contamination was checked by quantitative PCR (qPCR). The quality and quantity of treated RNA were analyzed using a DeNovix DS-11 spectrophotometer (DeNovix Inc.) and the Bioanalyzer 2100 system (Agilent Technologies) with an RNA integrity number ≥ 6 for cDNA library preparation. The rRNA depletion and Illumina libraries were made following the Illumina protocol (High-throughput sequencing facility of the I2BC, Gif sur Yvette, France). The *cDNA* samples were sequenced using Illumina NextSeq v. NS500446 (High-throughput sequencing facility of the I2BC, Gif sur Yvette, France), yielding, on average, 22.8 x 10^6^ 50 nt. paired-end reads (+/- 6.8 x10^6^ reads).

#### RNA-seq data analysis

Read sequences were mapped on our reference genome sequence with BWA mem [v. 0.7.9a-r786] using default settings, and coverage values of all genomic features were computed with the bedtools “coverage” command [v2.17.0] [73]. RNA differential gene expression analysis was performed with the DESeq R-package [v. 1.39.0] [74].

### Western-blot analysis

The protein extractions and Western-blot analyses were performed as previously described [20]. The membranes were incubated overnight at 4°C with a 1:5000 dilution of monoclonal rabbit anti-HA antibodies (Sigma-Aldrich) or 1:5000 dilution of monoclonal rabbit anti-FLAG antibodies (Sigma-Aldrich).

### Chromatin immunoprecipitation (ChIP)

Exponentially growing *D. radiodurans* cells (100 mL, A_650nm_ = 0.7) expressing DdrO-V5 or native DdrO protein were cross-linked with 1% formaldehyde in TGY2X medium for 25 min at 30°C with continuous shaking. Crosslinking reactions were quenched by the addition of 125 mM glycine for 15 min. Cells were harvested by centrifugation (4000 *g*, 10 min, 4°C), washed twice with cold Tris Buffer Solution (TBS, 50 mM Tris, 100 mM NaCl pH 7.5) and then resuspended in 3 mL of lysis buffer (160 nM NaCl, 20 mM, Tris-HCl pH7.5, 1 mM EDTA, protease inhibitor cocktail (Roche)). Cells were disrupted and DNA sheared using a One Shot Cell Disruptor (CellD SARL) to an average size of 100-300 bp (2 rounds of 2.4 kbar). Insoluble material was removed by centrifugation at 20,*000 g* for 10 min at 4°C and the supernatant was collected in a sterile microcentrifuge tube. Then, 500 µL of supernatant fluid was added to 25 µL of pre-incubated protein G magnetic beads (ChIP-Adembeads ChIP-Adem-Kit, Ademtech SA) with 5 µg of anti-V5 rabbit polyclonal antibody (ab9116, Abcam) in IP buffer (50 mM HEPES-KOH pH 7.5, 150 mM NaCl, 1 mM EDTA, 1% Triton 100, protease inhibitor cocktail (Roche)). After overnight incubation at 4°C with rotation, the immunoprecipitates were washed 5 times with washing buffers (ChIP-Adembeads ChIP-Adem-Kit, Ademtech SA).

Immune complexes were eluted in 200 µL of elution buffer. The eluted samples (20µL) were saved for control Western blots, and the remainder was incubated for 2 h at 37°C with shaking with 100 µg/mL Proteinase K. Then, the supernatant was incubated overnight at 65°C to reverse cross-linking with 100 µg/mL RNAse A. The DNA was purified using the PCR Clean-up kit (Macherey-Nagel).

### ChIP-seq

Raw FASTQ files were obtained from sequencing “IP”, “Mock” and “Input” samples comprising 19 and 13, 10^6^ sequences, respectively. The quality score was verified with FASTQC software (https://www.bioinformatics.babraham.ac.uk/projects/fastqc/) and Illumina adaptor sequences were removed with Cutadapt software (https://cutadapt.readthedocs.io/en/stable/). Sequence alignments on the genomic sequence were performed with Bowtie2 software (http://bowtie-bio.sourceforge.net/bowtie2/index.shtml) (default parameters). Output SAM files were converted and indexed into BAM files, using the Samtools software (http://www.htslib.org/). They were used both for visualization with IGV, and additional conversion into BED files with Bedtools software (https://bedtools.readthedocs.io/en/latest/index.html<u>) providing</u> the input file format required by bPeaks programs (https://cran.r-project.org/web/packages/bPeaks/index.html) to perform peak calling. Search for conserved motifs were performed by MEME and FIMO (https://meme-suite.org/meme/) with a Match pValue < 1.0 x 10^-4^. Prediction of *E. coli*-like gene promoter elements and transcription start site in gene promoters was carried out using BPROM (http://www.fruitfly.org/seq_tools/promoter.html) [75]. In order to sort data from Chip-seq and RNA seq and to integrate them with the conserved motifs found by MEME and FIMO, we used an in-house script (S1 File) which defines without *a priori* different lists of candidate genes, to be DdrO targets and hence provides detailed information about the process which was applied to obtain the results presented in the main text and Supplementary data files.

### Western blot analysis of RDR tagged-proteins

Exponentially growing of bacteria (15 mL, A_650nm_ = 0.3), grown at 30°C, were exposed to 1 or 5 µg/mL mitomycin C. After 3 h at 30°C with continuous shaking, cells were harvested by centrifugation at 4°C and the pellets washed with 1X cold saline-sodium citrate (SSC) buffer. Then, the bacteria pellets were re-suspended in 150 µL of SSC 1X with 0,4 mM protease inhibitor cocktail (Roche) and cells disrupted with a FastPrep Instrument using 0.1g of glass beads (500µm) and four pulses of 30 s. Cell debris were removed by centrifugation at 20,000 *g* for 10 min at 4°C and the supernatant fluid collected and placed in sterile microcentrifuge tubes. Protein concentrations were determined by Bradford assay (Bio-Rad). Proteins were subjected to electrophoresis through a 12% Glycine SDS-PAGE gel (Mini-PROTEAN TGX Stain-Free Precast gel, Bio-Rad) and transferred onto a polyvinylidene difluoride (PVDF) membrane (GE Healthcare). The Western-blot analyses was performed as previously described [20] with a 15,000 dilution of anti-V5 rabbit primary antibody (Abcam) or with a 1:5000 dilution of monoclonal rabbit anti-HA antibodies (Sigma-Aldrich).

### Sensitivity assay to DNA-damaging agents mitomycin C and UVC

Bacteria were grown in TGY2x liquid medium at 30°C to an A_650nm_ = 1 and sequential dilutions of cells were spotted on TGY plates supplemented (or not) with mitomycin C (60 ng/mL and 80 ng/mL at final concentration), exposed (or not) to UVC at a dose rate of 3.5 J/m^2^/s.

### Data reporting

The complete sequence and annotation of the genome have been deposited with the Genbank under accession number CP068791, CP068792, CP068793, CP068794. The complete high-throughput sequence data have been deposited with the Gene Expression Omnibus (GEO) data bank under accession number GSE175875 (RNA-seq and ChIP-seq).

## Acknowledgments

We acknowledge the High-throughput sequencing facility of the I2BC for its sequencing expertise and Pr Sébastien Bloyer for valuable discussions about ChIP-seq experiment. We thank M. DuBow for the English proofreading of the manuscript.

## Supplementary tables legends

**S1 Table.** List of CDS in the *D. radiodurans* strain ATC 13939 genome sequence. The paralogs found in this sequence as well as orthologs in other *D. radiodurans* R1 sequence releases are indicated with a threshold 80 % of maximum bit score applied.

**S2 Table**. Data integration for all CDS. List of CDS (Column A). Predicted RDR genes by Makarova et al [33] (Column B). RNA-seq data (columns C-F): Number of time points with differential expression in W37 (column C) or in D37 (column E), or only upregulated at intermediate time points, *i.e.* 6 h, 8 h and 16 h in W37 (column D) or in D37 (column F). ChIP-seq data (Columns G-L). Number of peaks detected in the promoter of each CDS (Column G), peak name (Column H) and their coordinates on Genome Orsay (Columns K-L). In case of the presence of two peaks all the information for the second peak is given in columns Y-AF. Search for palindromic or non-palindromic motifs with MEME or/and FIMO with an occurrence of one motif per sequence or any number of repetitions (Columns M-X). The sequences of each motif closed to the RDRM is indicated for each peak. For palindromic motifs, only the sequence found on one DNA strand is indicated.

**S3 Table**. Number of unique and common up- and downregulated genes in W37 and D37, respectively during the time course. For each time lapse, a 2-fold change expression threshold for the ratio experiments was applied together with a *p-Value* <0.01.

**S4 Table**. Variation of all CDS expression during the time course of W37 and D37 cells as well as *ddrO* and *spr* genes cloned in the *repU_T_*_s_ or the *repU*^+^ replication vectors, comparing each time point of the kinetic (4 h, 6 h, 8 h, 16 h and 24 h) to the first one (1 h). Red denotes upregulated genes (FC ≥ 2, *pVal* ≤ 0.01) and green denotes downregulated genes (FC ≤ -2, *pVal* ≤ 0.01). Genes that were previously predicted by Makarova [33] are underlined in yellow.

**S5-S7 Tables**. List of all the deregulated genes (S5 Table), upregulated (S6 Table) or downregulated (S7 Table) as differentially expressed in W37 strain between time lapses 1 h-6 h and 1 h-16 h in less than two comparisons (DE ≤ 2), and in more than three comparisons in D37 strain (DE > 3).

**S8-S11 Tables**. List of genes belonging to the RDR regulon in *D. radiodurans* (S8 Table) or matching two criteria only *i.e.* 6 genes with ChIP-seq and Transcriptomics criteria (S9 Table), 16 genes with Transcriptomics and RDRM criteria (S10 Table) and 25 genes with ChIP-seq and RDRM criteria (S11 Table). The coordinates of the ChIP-seq peaks as well as the genes regulated by DdrO are indicated for each replicon (Dro on Chr I, Dro_A: Chr II, Dro_C: plasmid). For RNA-seq assays, genes expression ratio in D37 and W37 species were reported for each time lapse using time 1 h as a reference. Red denotes upregulated genes (FC ≥ 2, *pVal* ≤ 0.01) and green denotes downregulated genes (FC ≤ -2, *pVal* ≤ 0.01). The predicted position of *E. coli*-like −35 and −10 like consensus sequences was carried out by BPROM [75] as well as the sequence and the position of the RDRM are pointed out from the predicted start of translation. (1-5) RDR genes previously predicted by bioinformatic analyses [33], by EMSA assays [26] or shown to be regulated after exposure to radiation by transcriptomic [12] proteomic studies [41] or in a Δ*irrE* mutant [24], respectively.

